# SpatialDDLS: An R package to deconvolute spatial transcriptomics data using neural networks

**DOI:** 10.1101/2023.08.31.555677

**Authors:** Diego Mañanes, Inés Rivero-García, Daniel Jimenez-Carretero, Miguel Torres, David Sancho, Carlos Torroja, Fátima Sánchez-Cabo

**Affiliations:** Centro Nacional de Investigaciones Cardiovasculares Carlos III (CNIC), 28029 Madrid, Spain; Departamento de Ingeniería Biomédica, ETSI de Telecomunicaciones, Universidad Politécnica de Madrid, 28040 Madrid, Spain

## Abstract

**Summary:** Spatial transcriptomics has changed our way to study tissue structure and cellular organization. However, there are still limitations in its resolution, and most available plaXorms do not reach a single cell resolution. To address this issue, we introduce SpatialDDLS, a fast neural network-based algorithm for cell type deconvolution of spatial transcriptomics data. SpatialDDLS leverages single-cell RNA sequencing (scRNA-seq) data to simulate mixed transcriptional profiles with predefined cellular composition, which are subsequently used to train a fully-connected neural network to uncover cell type diversity within each spot. By comparing it with two state-of-the-art spatial deconvolution methods, we demonstrate that SpatialDDLS is an accurate and faster alternative to the available state-of-the art tools.

**Availability and implementation:** The R package SpatialDDLS is available via CRAN-The Comprehensive R Archive Network: https://CRAN.R-project.org/package=SpatialDDLS. A detailed manual of the main functionalities implemented in the package can be found at https://diegommcc.github.io/SpatialDDLS.

**Supplementary information:** Supplementary data are available at Bioinformatics online.

## Introduction

Single-cell omics have represented one of the main technological advances towards the understanding of physiological and pathological states. However, the spatial context and localization of the cells are key elements with functional relevance that are missing with these techniques. In the last few years, spatial transcriptomics have revolutionized our ability to investigate biological processes by providing an unbiased way to understand tissue structure, cellular interaction, and function. Rather than studying cells as isolated and independent entities, it incorporates context through the spatial dimension while preserving the powerful information provided by whole transcriptome sequencing. However, due to the limitations of most available techniques, which fail to achieve single-cell resolution, computational methods are needed to identify the precise combination of cells within each spot. Deconvolution methods have been previously applied to bulk RNA-seq data in order to disentangle the cellular composition of samples from whole tissues or organs (Avila Cobos et al., 2018). For example, being able to quantify the different types of infiltrated lymphocytes in a given tumor starting from RNA-seq of the whole sample can serve as a very accurate method to predict the time-to-death from colorectal or breast cancer patients (Torroja and Sanchez-Cabo, 2019). A natural extension of these methods is to apply them to deconvolute the transcriptomics data from each sequenced spot in spatial transcriptomics to identify their exact cellular composition. There is a broad spectrum of tools which follow different approaches to solve this problem (Li et al., 2022), but most utilize single-cell RNA-seq (scRNA-seq) datasets from the same biological context as references, thereby presenting the issue as a supervised task. However, they usually rely on predefined markers defined either manually or through differential expression analysis, and typically have long running times that pose challenges for their practical application (Li et al., 2022).

In this work, we introduce SpatialDDLS, an R package that provides a fast neural network-based solution for cell type deconvolution of spatial transcriptomics data. The algorithm employs scRNA-seq data to simulate mixed transcriptional profiles with known cell composition, with which a fully-connected neural network is trained with the aim to uncover cell type diversity within each spot (Figure 1). In contrast to other methods which are computationally intensive and rely on a predefined and biased set of cell type markers, SpatialDDLS does not require the definition of cell identity signatures and has a lightweight computational processing. To demonstrate its performance and efficiency, we have benchmarked our tool against two state-of-the-art spatial deconvolution methods in two different biological contexts: murine lymph nodes upon stimulation (Lopez et al., 2022) and murine hippocampus samples (10x-Genomics, 2020; Saunders et al., 2018). SpatialDDLS’ predictions reproduced known cell type location paaerns and yielded similar results compared to other methods, while reducing the computational times by 1.8-14 fold, thereby making it a competitive alternative to already available tools.

**Figure 1:**
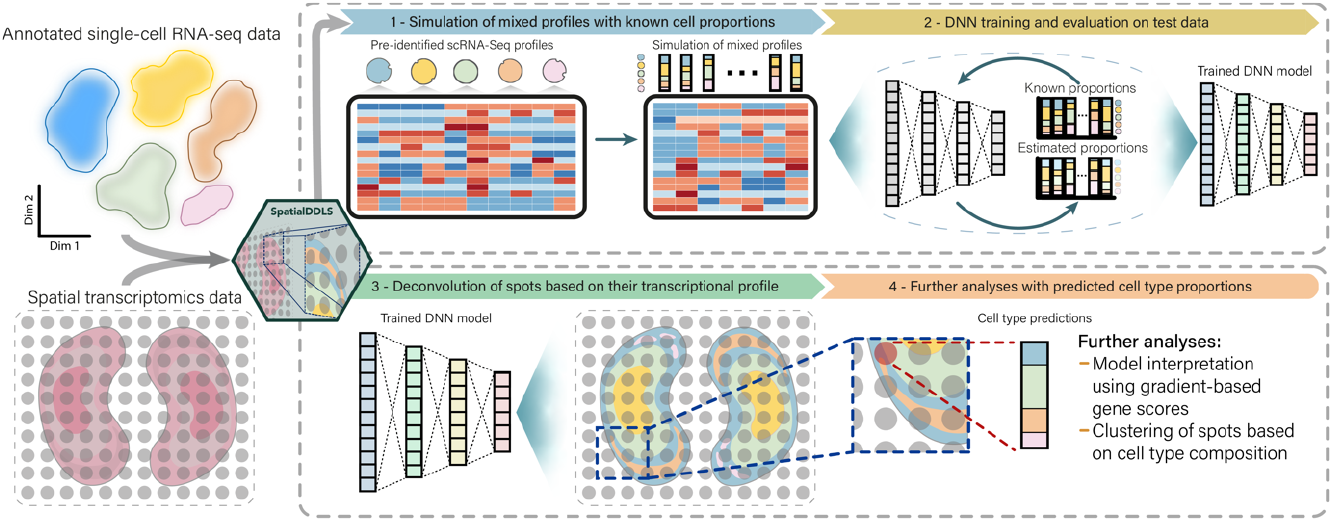
Schematic overview of SpatialDDLS. SpatialDDLS takes both an annotated single-cell RNA-seq dataset to be used as reference, and the Spatial transcriptomics datasets wanted to be deconvoluted. Then, it simulates mixed transcriptional profiles with known cell composition and trains a fully-connected neural network able to make accurate predictions of cell type proportions. These cell type proportions and the trained model can be used for further analyses.

## SpatialDDLS

SpatialDDLS is an extension of our deconvolution tool for bulk RNA-seq (Torroja and Sanchez-Cabo, 2019) implemented in the R package digitalDLSorteR (Mañanes et al., 2022). The algorithm uses scRNA-seq to simulate mixed transcriptional profiles for training neural network models capable of estimating the cell proportions of new mixed transcriptional profiles typically present in spatial transcriptomics data. It consists of three main steps (Figure 1):

1. Simulation of mixed transcriptional profiles with known cellular proportions. SpatialDDLS begins by using a pre-identified scRNA-seq dataset that is partitioned into training and test cell subsets. Then, cell proportions are simulated from each labelled subset of cells by using different approaches (see Supplementary Methods). Once the cellular composition matrix is generated, training and test mixed transcriptional profiles with those cell proportions are simulated.
2. Neural network training and evaluation. A neural network model is trained and evaluated using the simulated mixed profiles. Thanks to the inclusion of a test subset, this workflow allows for an assessment of whether the model is correctly identifying the transcriptional features of every cell type considered in the reference.
3. Deconvolution of spatial transcriptomics datasets. Once the model is trained, it is capable of making whole transcriptome-based predictions of cell type proportions in new samples, enabling the deconvolution of whole spatial transcriptomics datasets.

All these steps are implemented using the S4 object-oriented programming system of R to centralize all intermediate data generated during the workflow and provide a user-friendly usage. Regarding its implementation, SpatialDDLS makes use of the keras (Allaire and Chollet, 2021) and tensorflow (Allaire and Tang, 2021) R packages for all deep learning-related tasks, and S4-classes from the Bioconductor’s environment (Huber et al., 2015) for the storage of gene expression matrices (scRNA-seq and spatial transcriptomics). Therefore, it can be entirely integrated into the typical workflow used for the analysis of transcriptomics data in R. In addition, the possibility to work with The Hierarchical Data Format version 5 (HDF5) files as back-end has been implemented at each step of the workflow by using the DelayedArray (Pagès, 2021a) and HDF5Array (Pagès, 2021b) R packages to provide a way to handle large amounts of data on RAM-constrained machines. For a detailed explanation of each step with code and examples, see the website of the package (https://diegommcc.github.io/SpatialDDLS).

## Results

To assess the performance of SpatialDDLS, we decided to analyze samples from two different biological contexts that exhibit clear spatial paaerns in their cell type distribution: mouse hippocampus (Figure S1a) (Saunders et al., 2018) and mouse lymph node (Figure S3a) (Lopez et al., 2022) samples.

Firstly, we evaluated the model training and performance by using simulated mixed transcriptional profiles from these two experiments. As shown in Figures S1b and S3b, the trained models achieved excellent performance on simulated data of both datasets (R ≥ 0.98 (Pearson’s correlation coefficient) and CCC ≥ 0.96 (concordance correlation coefficient) for all cell types), indicating that they effectively detected biological signals for every cell type.

Next, we benchmarked SpatialDDLS against two state-of-the-art methods in the spatial transcriptomics field: cell2location (Kleshchevnikov et al., 2022) and RCTD (Cable et al., 2022). We chose these tools because of their high performance in different recently published benchmarks (Li et al., 2022, 2023; Yan and Sun, 2023). In both experiments, SpatialDDLS generated similar predictions to those of cell2location and RCTD. This agreement was confirmed by hierarchical clustering of Pearson’s correlation coefficients between the estimations made by each method for each cell type, showing that they were mostly clustered by cell type rather than the method used (Figure S1c and S3c). Then, we plotted the estimations of the most representative cell types of each dataset together with the expression of classical markers (Table S1) to observe their spatial location (Figure S2 and S4). The three methods yielded similar spatial patterns that co-localized with the expression of their markers, although also some differences were observed for specific cell types. In particular, SpatialDDLS tended to predict higher proportions of endothelial cells and astrocytes in the hippocampus sample than cell2location, which is in line with the expression of their markers and RCTD predictions (Figure S2). In the lymph node dataset, CD8+ T cells were underestimated by SpatialDDLS, although CD4+ T cell and migratory dendritic cell signals were better captured compared with RCTD estimations (Figure S4). Overall, this demonstrates a high agreement between the three methods but the presence of specific tendencies, making their predictions complementary to each other.

Importantly, despite these similar results, SpatialDDLS exhibited a significant improvement in running time compared with the other two tools (Table S2). While cell2location showed the worst performance in both datasets, RCTD kept good running times in the hippocampus samples. However, when the number of spots considered increased as in the lymph node samples, the algorithm did not scale well compared with SpatialDDLS, which kept similar running times in both datasets.

## Conclusion

SpatialDDLS is a flexible spatial deconvolution tool of easy use fully integrated in the R/Bioconductor ecosystem. We have demonstrated that it generates comparable results to those of two state-of-the-art methods while keeps shorter running times. All these features make it a robust alternative to existing methods. In addition, SpatialDDLS does not need the definition of a set of markers for each cell type and performs whole-transcriptome predictions. We think that this fact can be useful in the context of paired scRNA-seq and spatial transcriptomics datasets, as SpatialDDLS could account for specific transcriptional features that cell types can undergo depending on the biological context.

## Acknowledgements

The authors thank the Bioinformatics Unit for the support and the insightful discussions about the project.

## Funding

This work was supported by the Ministerio de Ciencia, Innovación, y Universidades (MCIU) [grant no. RTI2018-102084-B-I00], by the Ministerio de Ciencia e Innovación MCIN/AEI/10.13039/501100011033 [grant no. TED2021-132296B-C54] and by the European Union NextGenerationEU/PRTR. The CNIC is supported by the Instituto de Salud Carlos III (ISCIII), the Ministerio de Ciencia e Innovación (MCIN) and the Pro CNIC Foundation, and is a Severo Ochoa Center of Excellence (grant CEX2020-001041-S funded by MCIN/AEI/10.13039/501100011033). DM is supported by a predoctoral grant from the Spanish Government (grant number PRE2020-092578 MCIN/AEI/10.13039/501100011033). IRG received the support of a fellowship from”la Caixa” Foundation (ID 100010434, fellowship code: LCF/BQ/DR20/11790019).

## Conflict of Interest

none declared.

## Supplementary Material

## Supplementary Methods

### SpatialDDLS overview

SpatialDDLS is an R package which provides a user-friendly approach to deconvoluting spatial transcriptomics data. The package offers several functionalities for effectively handling large datasets through the utilization of HDF5 files and streamlines the overall process with a few simple function calls. It tries to offer a framework in which deconvolution of spatial transcriptomics data using neural networks is easy and flexible depending on the particularities of each problem. The algorithm comprises three main steps.

#### 1 - Loading data

SpatialDDLS operates within a supervised framework, requiring thus both the spatial transcriptomics wanted to be deconvoluted and a reference single-cell RNA-seq (scRNA-seq) dataset with pre-identified cell types. The goal is to consider only those genes that are relevant in both types of data for further steps. To do so, it starts by applying two filters to reduce the number of original dimensions. These filters must be modified according to the particularities of the data (number of cells, sequencing depth, etc.):

1. Minimum count cutoff for N cells.
2. Minimum non-zero average count cutoff per cell type, only applied to the scRNA-seq.

Then, only genes shared between both modalities are retained. In addition, if multiple spatial transcriptomics slides are provided, SpatialDDLS offers the option to keep only those genes present in a specified number of slides. These steps aim to expedite subsequent steps by avoiding the consideration of the entire noisy expression matrix. Notably, when working with massive amounts of data, single-cell profiles can be provided as HDF5 files. SpatialDDLS handles this format by using the DelayedArray (Pagès, 2021a) and HDF5Array (Pagès, 2021b) R packages.

#### 2 - Simulation of mixed transcriptional profiles

The second step consists of the generation of mixed transcriptional profiles with known cell compositions. This is achieved by the genMixedCellProp function, which generates the cell composition matrix, and simMixedProfiles, which simulates the mixed profiles and saves them in the SpatialDDLS object. SpatialDDLS is again able to accommodate these simulated samples using HDF5 files as back-end, but this option is not necessary for most situations.

In the cell composition generation process, the package initially partitions the single-cell profiles into training and test subsets (default split ratio of 0.66). The cell proportions for each simulated profile are then determined using three different approaches:

##### 1. Random profiles

In this approach, profiles are generated by randomly sampling from a Dirichlet distribution. To introduce sparsity in the cell proportions, the prob.sparity parameter is used, controlling the probability of having missing cell types in each simulated profile instead of a mixture of all cell types.

##### 2. Pure profiles

This approach involves aggregating a specified number of cells from the same cell type to create pure profiles. These samples aim to generate less noisy representations for each cell type considered in the experiment, as opposed to directly using the single-cell profiles.

##### 3. Forced-sparse profiles

This approach enforces a minimum number of missing cell types, resulting in sparse samples (min.zero.prop parameter). Proportions for the remaining cell types are then generated using a Dirichlet distribution.

These methods are designed to introduce greater sparsity in terms of cell type composition, enabling the neural network to learn to detect scenarios where certain cell types may be missed. For analyses similar to those presented in this article, we recommend generating a total of 15,000-25,000 mixed transcriptional profiles (num.sim.spots). The percentage of profiles generated by each method can be controlled using the proportion.method parameter, but the default values are suitable for most situations. Each sample comprises a specific number of aggregated cells with a default of 50 single-cell profiles per sample. Finally, in cases where certain cell types are underrepresented, SpatialDDLS provides the option to simulate new single-cell profiles using the ZINB-WaVE framework (Risso et al., 2018) by the estimateZinbwaveParams and simSCProfiles functions. This feature enables the increase of cell type-specific signals, thereby increasing their representation through an augmentation-based approach.

In the simulation step, single-cell profiles can be aggregated using different strategies, but aggregating raw counts by summing them up is the preferred one. Then, mixed transcriptional profiles are normalized to account for sequencing depth using counts per million and log2-transformed.

#### 3 - Training the neural network model and deconvolution of spatial transcriptomics datsets

Once the SpatialDDLS object contains the normalized mixed transcriptional profiles, a neural network model is trained using the training subset. To address potential issues arising from variations in gene scales, the profiles are feature-wise rescaled between 0 and 1. While alternative transformations are available, this rescaling strategy is the default choice.

The package implements a default architecture for the neural network model, but users have the flexibility to fully customize it based on the specific characteristics of their datasets. However, we recommend considering the following parameters as a starting point, although they may vary depending on the complexity of the dataset being deconvoluted (number of cell types, similarity between them, etc.):

- Architecture: Two hidden layers with 300 neurons each.
- Activation function for hidden layers: Sigmoid function as the activation function.
- Optimization function: Kullback-Leibler divergence, which is suitable for modeling probability distributions.
- Batch size: 64 samples.
- Number of epochs: 60-80.
- Dropout regularization: One dropout layer with a rate of 25% to every hidden layer to prevent overfitting.
- Activation function for the last layer: Softmax function, enabling the neural network to predict probabilities that can be interpreted as cell type proportions.

After training, the model becomes capable of predicting the cell composition of new samples based solely on their transcriptional features. Additionally, the package also includes several functions for inspecting and evaluating the model’s predictive performance. These functions allow users to assess whether their models are effectively learning patterns associated with specific cell types, and thus determining if adjustments to the hyperparameters are required.

### scRNA-seq and spatial transcriptomics datasets

For the mouse hippocampus analysis, the scRNA-seq dataset used as reference is sourced from Saunders et al. 2018 and was downloaded from the Broad Institute’s Single Cell Portal (SCP948 study). The spatial transcriptomics dataset was obtained from the 10x Genomics website (10x-Genomics, 2020). In the case of the mouse lymph node analysis, the scRNA-seq and spatial transcriptomics data were obtained from Lopez et al. 2022 and are available at the Gene Expression Omnibus database under the accession number GSE173778.

Both datasets were analyzed and visualized using Seurat v4.1 (Hao et al., 2021). In brief, most variable genes were taken and used as input for principal component analysis (PCA). Then, first 15 principal components were used for visualization through Uniform Manifold Approximation and Projection for Dimension Reduction (UMAP) using default parameters implemented in Seurat.

### Deconvolution of spatial transcriptomics datasets

Unless stated, the parameters used for each model presented in this article are the ones the package has by default.

#### 1- Mouse hippocampus

Only genes shared between the single-cell reference and the spatial transcriptomics dataset were used with no further filtering, resulting in 273 genes. Then, 15,000 mixed transcriptional profiles were simulated and 66.6% of them used to train for 60 epochs a neural network of two hidden layers with 300 neurons each.

#### 2- Mouse lymph node

In the scRNA-seq dataset, genes were filtered setting a cutoff of three counts in five cells and a cutoff of non-negative average expression of three. Spatial transcriptomics dataset genes were filtered setting a cutoff of one count in one spot. Then, only shared genes between both modalities were considered for further analysis, resulting in a total of 1,300 genes. Then, 15,000 mixed transcriptional profiles were simulated and 66,6% of them used to train for 60 epochs a neural network model of two hidden layers with 300 neurons each.

### Comparison with cell2location and RCTD

For the analyses conducted using cell2location and RCTD, the default parameters and tutorial recommendations were used. Particularly:

- cell2location: We followed the tutorials available on the cell2location documentation website: https://cell2location.readthedocs.io. The single-cell regression model was trained with parameters max_epochs=250, lr=0.002. The cell2location model was obtained with parameters max_epochs=30,000. Then, cell2location’s predictions in each spot were divided by the maximum to treat them as cellular proportions.
- RCTD: We followed the tutorials present in the GitHub repository (https://github.com/dmcable/spacexr.git). The”full” mode was selected.

To compare the three methods, estimated cell proportions of each tool were analyzed by calculating the Pearson’s correlation coefficients of estimated cell proportions per method and cell type. Then, Pearson’s correlation coefficients were clustered using hierarchical clustering and plotted as a heatmap using the ComplexHeatmap R package (Gu et al., 2016). All deconvolution analyses (including SpatialDDLS) were performed with an Intel(R) Core(TM) i5-10500U CPU @ 3.80 GHz with 32 GB of RAM.

### Expression of classic markers

To determine if the predicted cell type proportions are consistent with the expression levels of established cell type markers, we computed the mean Z-score expression levels of a manually selected set of markers for each cell type (see Table S1). This analysis was performed primarily to visualize the spatial distribution of specific cell types and not for quantitative comparison purposes.

## Code availability

The source code for SpatialDDLS is available at https://github.com/diegommcc/SpatialDDLS, and it is also available on CRAN https://CRAN.R-project.org/package=SpatialDDLS.

## Supplementary Figures and Tables

**Figure S11:**
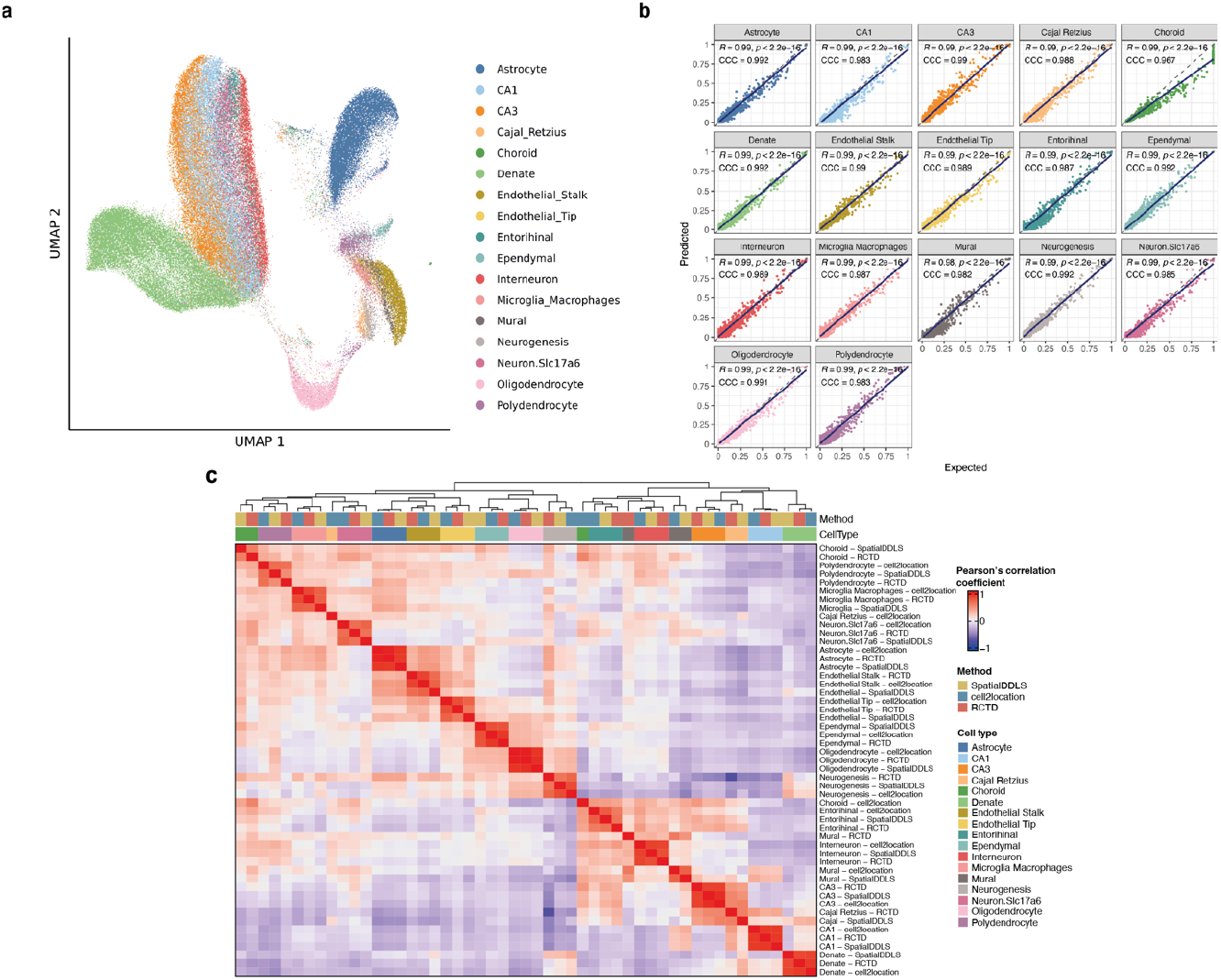
Mouse hippocampus dataset. **a**. UMAP representation of scRNA-seq data from Saunders et al., 2018. **b**. Correlation between expected and predicted cell proportions of simulated mixed transcriptional profiles for every cell type. **c**. Heatmap of Pearson’s correlation coefficients between cell type proportions estimated by each method in hippocampus spatial transcriptomics data (10x-Genomics, 2020). Abbreviations: CA: cornu ammonis.

**Figure S2:**
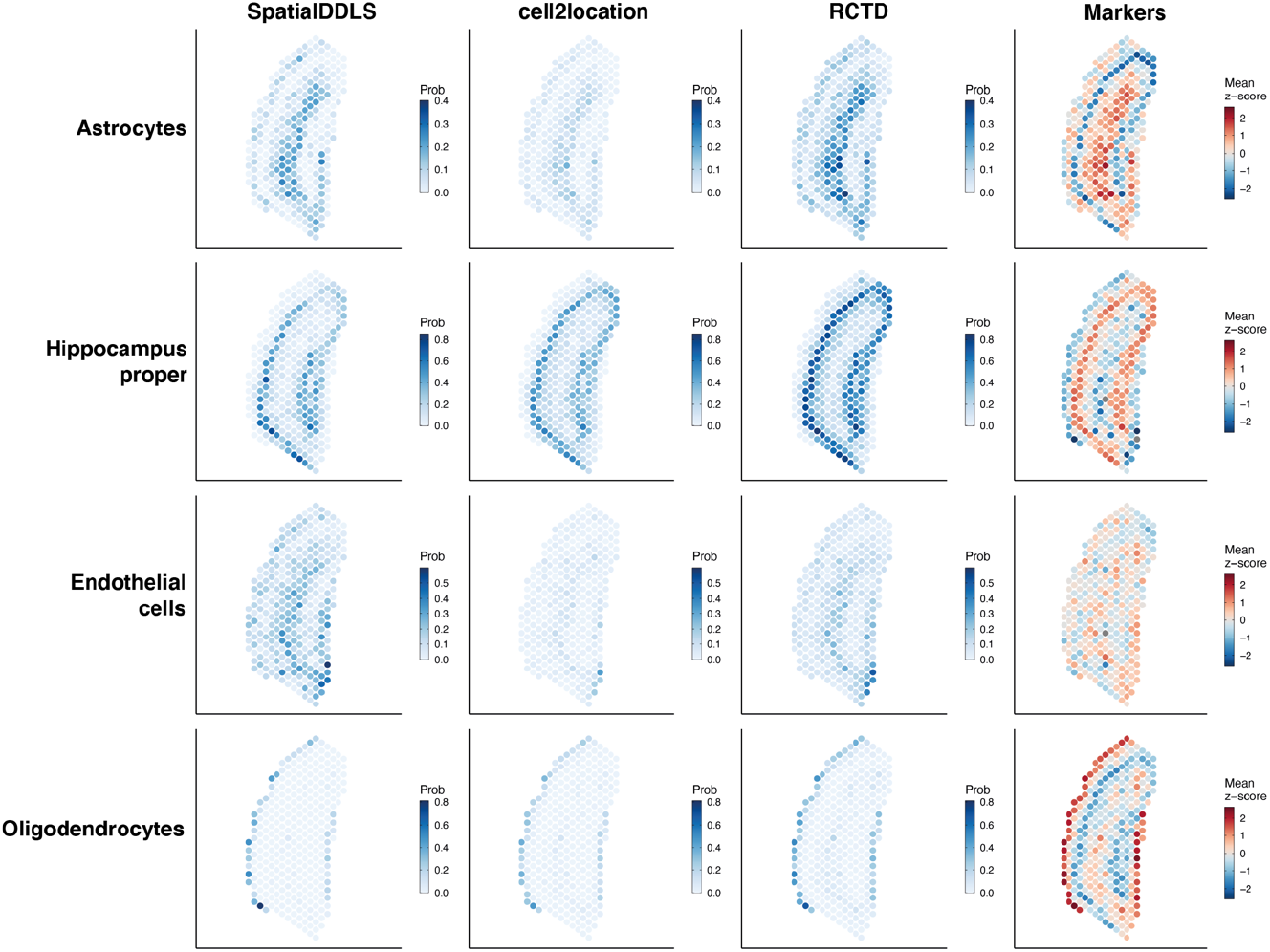
Estimated cell type proportions mapped to the spatial coordinates in the mouse hippocampus dataset. Endothelial cells and hippocampus proper were generated by adding up predicted proportions of: Endothelial cells = Endothelial stalk + Endothelial tip; Hippocampus proper = CA1 + CA3 + Dendate.

**Figure S3:**
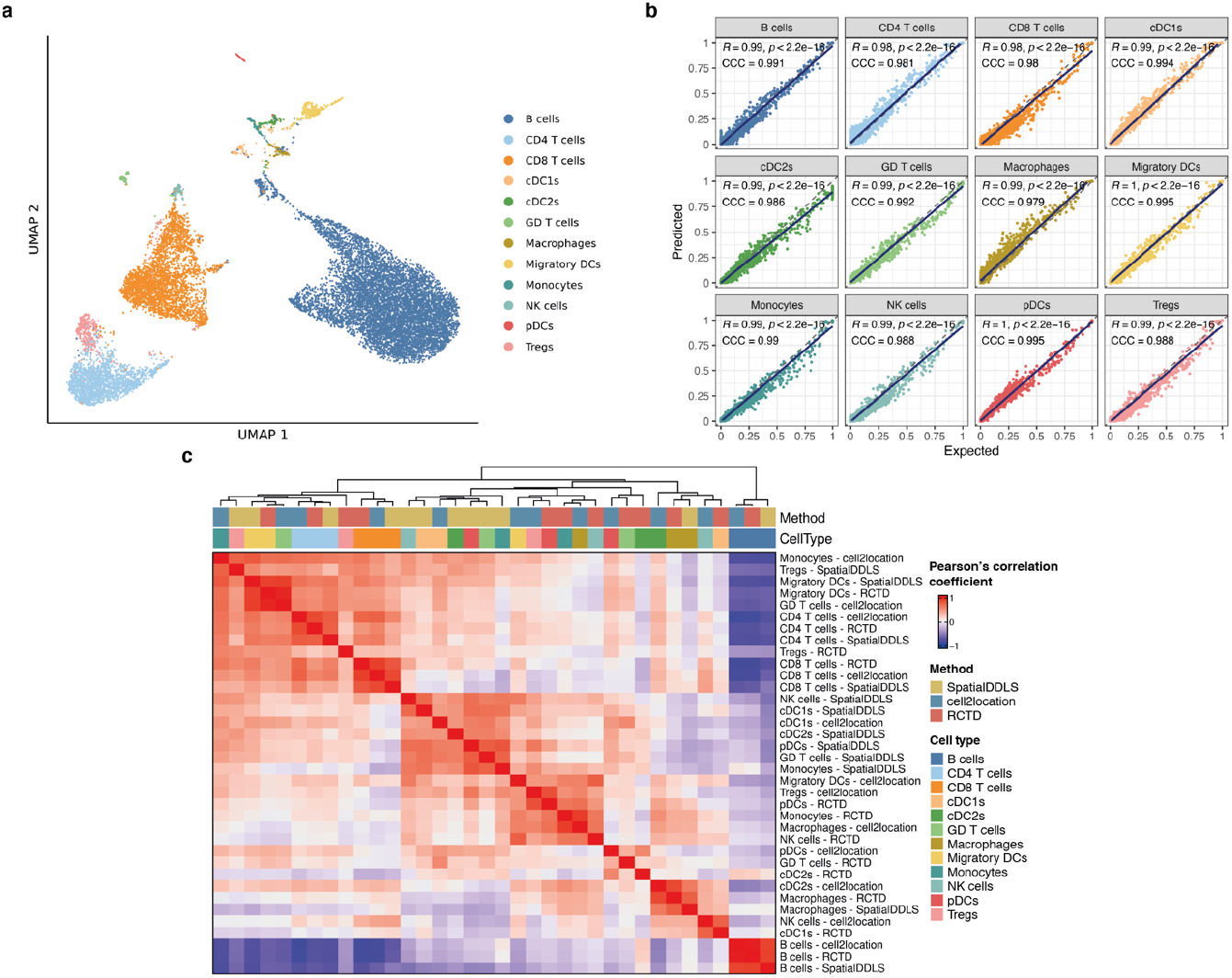
Mouse lymph node dataset. **a**. UMAP representation of scRNA-seq data from Lopez et al., 2022. **b**. Correlation between expected and predicted cell proportions of simulated mixed transcriptional profiles for every cell type. **c**. Heatmap of Pearson’s correlation coefficients between cell type proportions estimated by each method in lymph node spatial transcriptomics data (Lopez et al., 2022). Abbreviations: cDC: conventional dendritic cells; GD: gamma-delta; DCs: dendritic cells; pDCs: plasmacytoid dendritic cells; Tregs: T regulatory cells.

**Figure S4:**
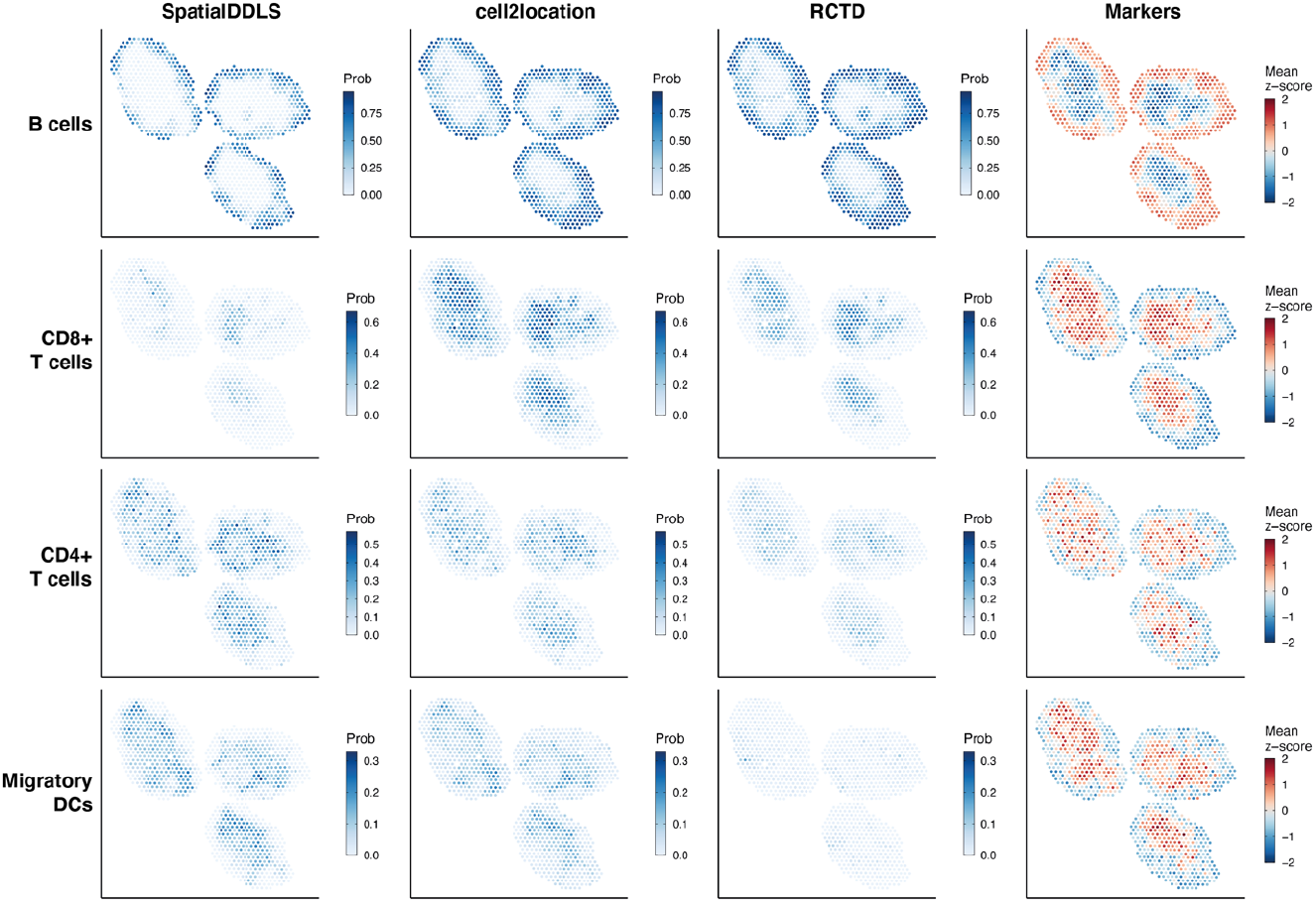
Estimated cell type proportions of representative cell types mapped to the spaptial coordinates in the mouse lymph node dataset. Abbreviations: DCs: dendritic cells.

**Table S1:** Cell type markers used to check spatial location of main cell types.

| Dataset | Cell type | Markers |
| --- | --- | --- |
| Hippocampus | Astrocytes | Slc1a2 Apoe Aldoc |
| Hippocampus | Hippocampus proper | Nrgn Ywhaz |
| Hippocampus | Endothelial cells | Sparcl1 Ptgsd Gpm6a |
| Hippocampus | Oligodendrocytes | Mbp Cnp Ptgsd |
| Lymph node | B cells | Cd74 Cd19 Cd79a Cd79b Ly6d |
| Lymph node | CD4+ T cells | Cd4 Lef1 Fyb |
| Lymph node | CD8+ T cells | Cd8b1 Cd8a Trac |
| Lymph node | Migratory DCs | Ccl5 Anxa3 Fscn1 |

**Table S2:**
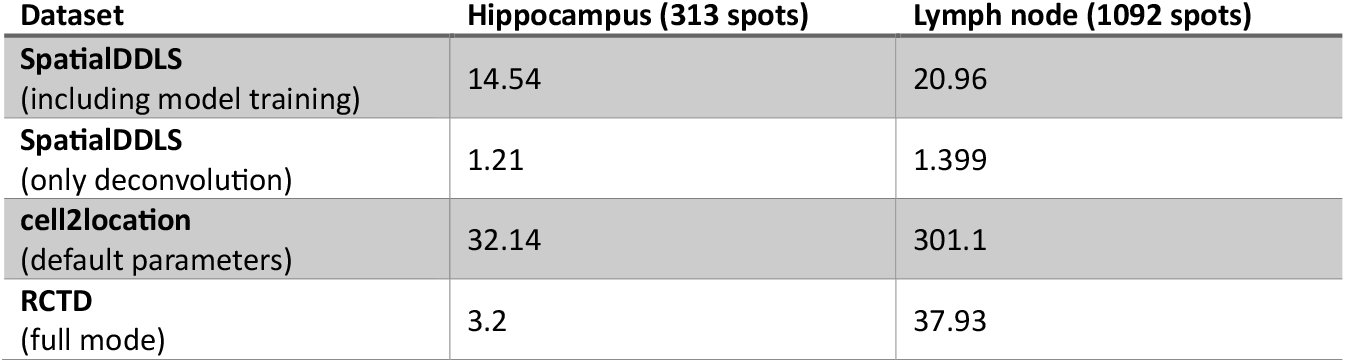
Running times (minutes) of benchmarked deconvolution algorithms on mouse hippocampus and lymph node samples.

## Notes

https://diegommcc.github.io/SpatialDDLS

### Competing Interest Statement

The authors have declared no competing interest.

